# Semantic Probing: Feasibility of using sequential probes to decode what is on a user’s mind

**DOI:** 10.1101/496844

**Authors:** Karen Dijkstra, Jason Farquhar, Peter Desain

## Abstract

In this paper, we investigate the feasibility of using multiple sequential probe words to decode their relatedness to an active semantic concept on a user’s mind from the respective electrophysiological brain responses. If feasible, this relatedness information could be used by a Brain Computer Interface to infer that semantic concept, by integrating the knowledge of the relationship between the multiple probe words and the ‘unknown’ target. Such a BCI can take advantage of the N400: an event related potential that is sensitive to semantic content of a stimulus in relation to an established semantic context. However, it is unknown whether the N400 is suited for the multiple probing paradigm we propose, as other intervening words might distract from the established context (i.e., the target word). We perform an experiment in which we present up to ten words after an initial target word, and find no attenuation of the strength of the N400 in grand average ERPs and no decrease in classification accuracy for probes occurring later in the sequences. These results lay the groundwork for the development of a BCI that infers the concept on a user’s mind through repeated probing.

## Introduction

Brain Computer Interfaces (BCIs) use brain activity as a direct input for a computer. For instance, many BCI applications are designed to offer alternative means of communication to people that are no longer able to use conventional input devices (e.g., keyboards). Existing communication BCIs come in a wide array of paradigms, achieving communication in different ways. These paradigms range from binary choice selection, allowing, for instance, the selection of “Yes” or “No” through various approaches: e.g., through the use of mu/beta rhythms^1^, auditory P300s^2^ or fNIRS^3^ — to spellers, in which characters can be selected to write a message: e.g., matrixspellers using the P300^4^, or Broad Band Evoked Potentials^5^, spellers using attention to tactile stimuli^6^ or visual spellers using rapid stimulus presentation (RSVP)^7^. Each have their strengths and weaknesses, allowing the choice of BCI to vary depending on the needs and preferences of a user.

Here, we are working towards a novel type of BCI that allows for the selection of words as a unit. In an approach similar to the game ‘20 questions’, in which someone tries to guess the person or object that someone is thinking of by asking only yes-no questions, we envision a BCI that infers the concept on a user’s mind by, in essence, asking them if a presented word is related to the to-be-inferred concept. More specifically, the proposed BCI presents a probe word, uses the electrophysiological responses of semantic relatedness to infer the probe’s relation to the target concept, and updates its belief state based on this evidence. It then presents a new probe, measuring another brain response, and so on, until it has sufficient confidence that the target concept has been identified. This can be employed as a word selection application, for purposes of communication (e.g., in sentence generation, or perhaps topic selection) as an alternative for existing BCI communication approaches, or for tip-of-the-tongue scenarios, in which the word for a to-be-communicated concept cannot be retrieved, which can be a problem for patients with anomic aphasia^8^, for instance.

Such an approach toward word selection has been outlined in the past^9^. In that study, users encoded a probe’s relatedness status using deliberate responses: Users were presented with a stream of probes and asked to press a button when a presented word was related; a task designed to simultaneously induce a movement related de-synchronization and elicit a P300 for related probes.

While using such deliberate signals is one way to approach such a BCI, there already exists a brain response that is inherently sensitive to the semantic content of a stimulus: the N400. It is an Event Related Potential (ERP) characterised as a wave that is more negative when a presented stimulus is unrelated, compared to a related stimulus, in relation to a previously established context^10^. Over the years it has proven a robust effect that can be elicited in a sentence context, in which the response to a word is measured based on its relation to earlier parts of the sentence, or in a word-pair context, in which a prime is presented followed by a related or unrelated probe. It is not only elicited with text on a screen, but also occurs when stimuli are pictures^11^ or when presented auditorily, as speech^12,13^, though the N400’s scalp distribution can vary. Importantly, it is not necessary that subjects are explicitly tasked to evaluate the semantic content of a stimulus to elicit an N400, though it appears to require that a stimulus’ meaning is processed (i.e., in a task where only the length of the word is relevant, no semantic priming effects from previous stimuli occur;^14,15^).

Using such an “automatic” brain response might be more pleasant for the user, as they have to provide less effort in deliberately evoking the right response. However, if we want to use the N400 for this BCI paradigm, we first have to establish whether this signal lends itself to such an approach.

First, the N400 should be detectable from a brain response to a single stimulus presentation, and not merely be observable in grand averages across or within participants. Previous work has shown that indeed the N400 can be detected from single word pair presentations (i.e., a target word and a single probe), with classification rates across individuals ranging from 54% to 67%^16^.

Secondly, to infer the concept on a user’s mind, a system would have to present a number of probe words consecutively, for the same concept, in order to collect sufficient evidence. Such a sequence of probe words could conceivably influence the context on a users mind, and it is therefore possible that results from a prime-single probe paradigm do not generalise to a setup with multiple (i.e., many) probes.

In other words, we want to know whether single trial classification of relatedness based on the N400 is still possible when multiple consecutive probes are presented following an initial target word. To establish what the effects of multiple consecutive probes are on the decoding of semantic relatedness, we designed an experiment in which we present a target word for participants to actively keep in mind, followed by up to ten probe words that are either related or unrelated to this target. While presenting ten probes is not necessarily sufficient for decoding purposes, we will use this data to determine if there is any indication of attenuation (i.e., a reduction of the magnitude) of the N400, when comparing first probes and probes late in the sequence. Specifically, we will compare grand average ERPs in response to probes appearing immediately after the target, with those in response to probes at the end of the sequence, combining data across participants to increase the sensitivity for detecting a difference.

In addition to this, there is some evidence that tasks designed to elicit N400s can also induce changes in oscillatory brain responses^17^. To evaluate this we will analyse the data in the frequency domain to determine if there are any spectral differences that could be used as additional features for decoding semantic relatedness.

## Results

### Behavioural results

In each trial, participants were presented with a target word to keep in mind, followed by up to ten probe words. At the end of each trial, participants indicated with a button-press whether the most recently presented probe had been related or unrelated to the initial target word (left vs. right button-press respectively). We compared the answers of each participant to the label of that probe and calculated the percentage of agreeing answers across all trials. Agreement between the response and label ranged from 70% to 95% across participants (mean 87%). A mismatch between the response and label can reflect a user error, or a disagreement on the relatedness status of the probe, as relatedness judgements can vary between individuals. Average reaction times per participant ranged from 368 ms to 1420 ms (mean 632 ms), with a strong correlation between reaction time and age (r = 0.80, n = 20, p = 0.00003). On average, reaction times to related probes were somewhat faster than to unrelated probes (578 ms and 650 ms respectively; agreeing responses only).

### Initial ERP analysis

To determine if there were any differences in brain responses to related compared to unrelated probes, we computed a grand average ERP, averaging across all related and unrelated probes presented in the experiment. This analysis produced an unexpected result: the grand average showed a difference in response between related and unrelated probes *prior to* stimulus onset, with the ERP for unrelated probes being more negative before, at, and some period after time 0. A difference in ERPs occurring prior to presentation of the stimulus of interest cannot reflect a brain response to that stimulus, suggesting that there is some structural factor (e.g., a previous stimulus) affecting the responses, that did not average out in the ERP. We hypothesise that this is a consequence of our stimuli design: while two related stimuli were never presented sequentially, unrelated stimuli could appear after either a related or an unrelated probe. If we split the the data into these three categories and compute grand averages we obtain the results in Figure 1a (for electrode Cz; baselined to the predecessor stimulus). Here we see that the pre-stimulus difference occurs only for unrelated probes occurring after a related probe.

**Figure 1.**
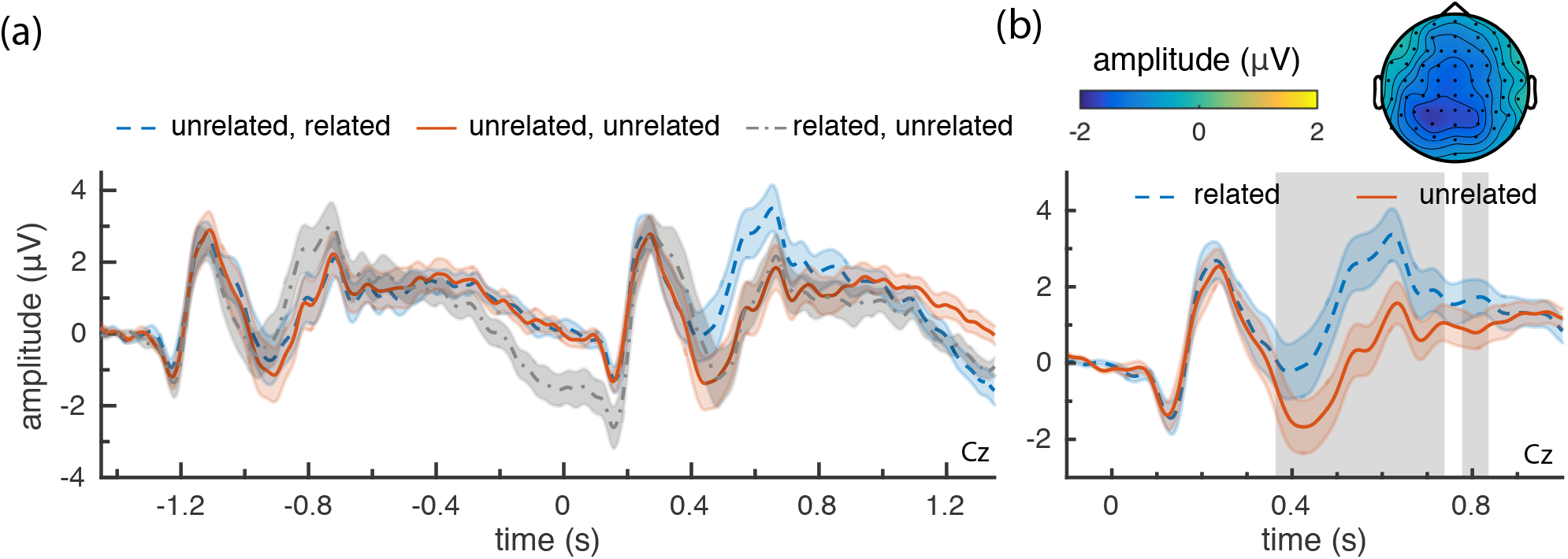
Grand average ERPs of responses to related and unrelated probes (Cz). (**a**) Responses to probes differentiated by their predecessor (related or unrelated; ‘*related, related*’ did not occur in the experiment) from −1.35 s (presentation of previous probe) to 1.35 s (presentation of next probe). (**b**) Grand average of related and unrelated probes that followed an unrelated probe. The grey area marks, for this channel, the timepoints that were part of the significant cluster identified by the cluster permutation test. The accompanying topoplot from 300 to 800 ms has been included on top (unrelated-related).

More specifically, there appears to be a late response that differs in amplitude between related and unrelated probes, which is still present when the next stimulus is presented. This is followed by a stimulus response (around 200 ms) after which the *related→unrelated* category behaves similarly to the *unrelated→unrelated*, until at least 1 s after stimulus presentation.

With responses to probes preceded by an unrelated probe appearing unconfounded (i.e., behaving more or less identically at stimulus onset), from here on we consider only these probe’s data, excluding the *related→unrelated* data (unless noted otherwise), to ensure that effects are not an artifact of our experimental design.

### ERP analyses

The resulting grand average ERP can be found in Figure 1b. Here we observe a difference between the two conditions from around 400 ms to 800 ms, where the response to unrelated probes is more negative than to related probes, as would be expected for an N400. This difference corresponds to the significant cluster found in a cluster permutation test^18^ (p = 0.001, two-tailed test, *α* = 0.05, n = 20, cluster marked in grey), confirming that there is a significant difference in the response to related and unrelated probes.

In order to determine whether or not the magnitude of the ERP decreases after more probes have been presented, we compare the ERP in response to the first probe in each plot, to the ERPs of those in final positions (i.e., the 9^*th*^ and 10^*th*^ position). Note, that due to the exclusion of unrelated probes that were preceded by a related probe, there are fewer instances available for unrelated probes in the final positions (104 in the first and 53 in the final position respectively, per subject). For both locations (first vs final), we plot the difference ERP (unrelated-related) in Figure 2a.

**Figure 2.**
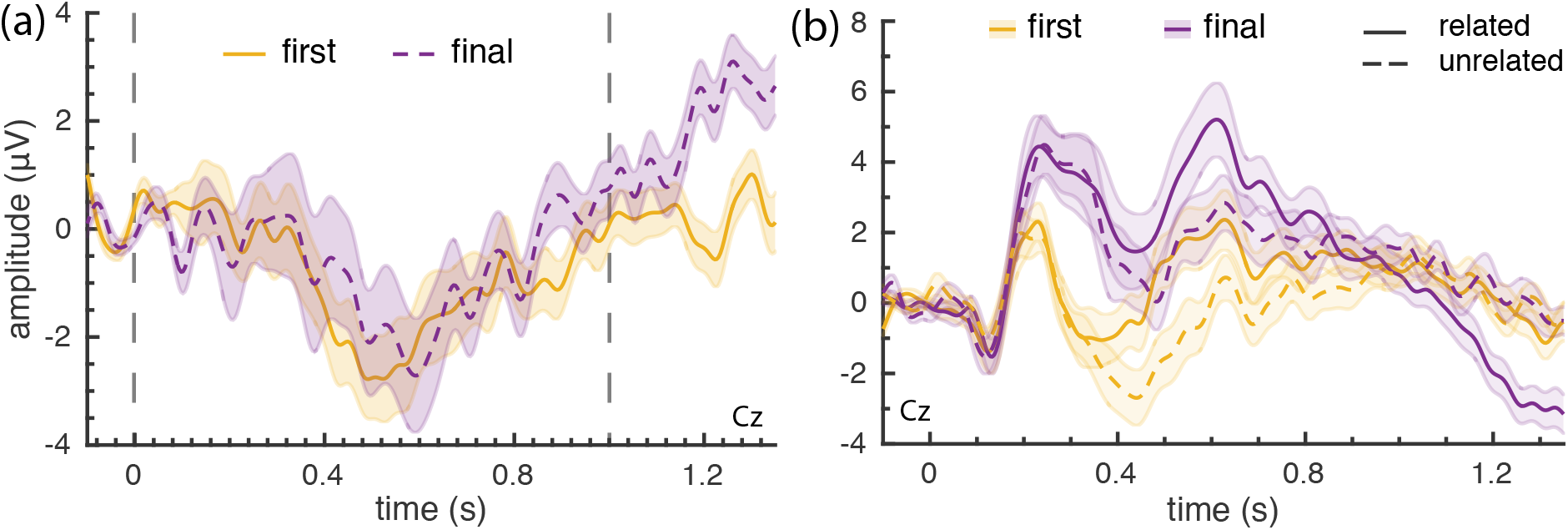
Grand average ERPs for probes presented immediately after a target (first; 1^*st*^ position), and probes presented at the end of a trial sequence (final; 9^*th*^ and 10^*th*^ position) (Cz). (**a**) Difference waves of probe responses: unrelated minus related. No significant clusters were found in a cluster permutation test applied to the period from 0 to 1 s (indicated by the gray frame). (**b**) Related and unrelated ERPs from first and final probe positions.

For both probes in the first and final positions, the response to unrelated probes was more negative than the response to related probes, resulting in a negative difference from around 400 ms to 800 ms. In the cluster permutation applied to the two difference waves (from 0 - 1 s; indicated by the grey frame), no significant clusters were found (lowest cluster p-value observed: p = 0.21, two-tailed test, *α* = 0.05, n = 20). Unexpectedly, the figure suggests that there is a difference after one second. To look at this in more detail, we plot the individual ERPs for related and unrelated for both the first and final positions in Figure 2b. Here, we see that ERPs in response to first and final probes look different, even though their difference waves are similar. Furthermore, the positive difference wave after 1 s in the final position, appears to be due to a late negative response for related probes in particular. This may be related to the late negative wave we observed in Figure 1a for related probes, which would imply this late negative wave did not occur for probes in the first position.

### Predicting relatedness

As our interest is ultimately in the decoding of these signals from single probe presentations, we trained classifiers to predict for a given probe response whether it is related or unrelated, for each subject, using a leave-one-sequence-out crossvalidation approach. No classifier was trained for subject 19, as they were unable to complete the full experiment. The accuracy per subject can be found in Figure 3a. Here, the accuracy of predicting whether a presented probe was related or unrelated is plotted per subject. In this figure, we shade the 99.75% binomial confidence interval of chance performance in red to get an indication of which participant’s results did not differ from chance (i.e., a Bonferroni-corrected 95% confidence interval around 50%). For 12 out of 19 participants, the classification accuracy could be distinguished from chance level. On average, the relatedness of probes was predicted correctly for 58.3% of probes (sd = 6.6%). To ensure our crossvalidation approach did not overestimate the classification accuracies, we compare them to the accuracies obtained with a train-test split (80%-20%). For this train-test split, the average accuracy across participants was 59.5%.

**Figure 3.**
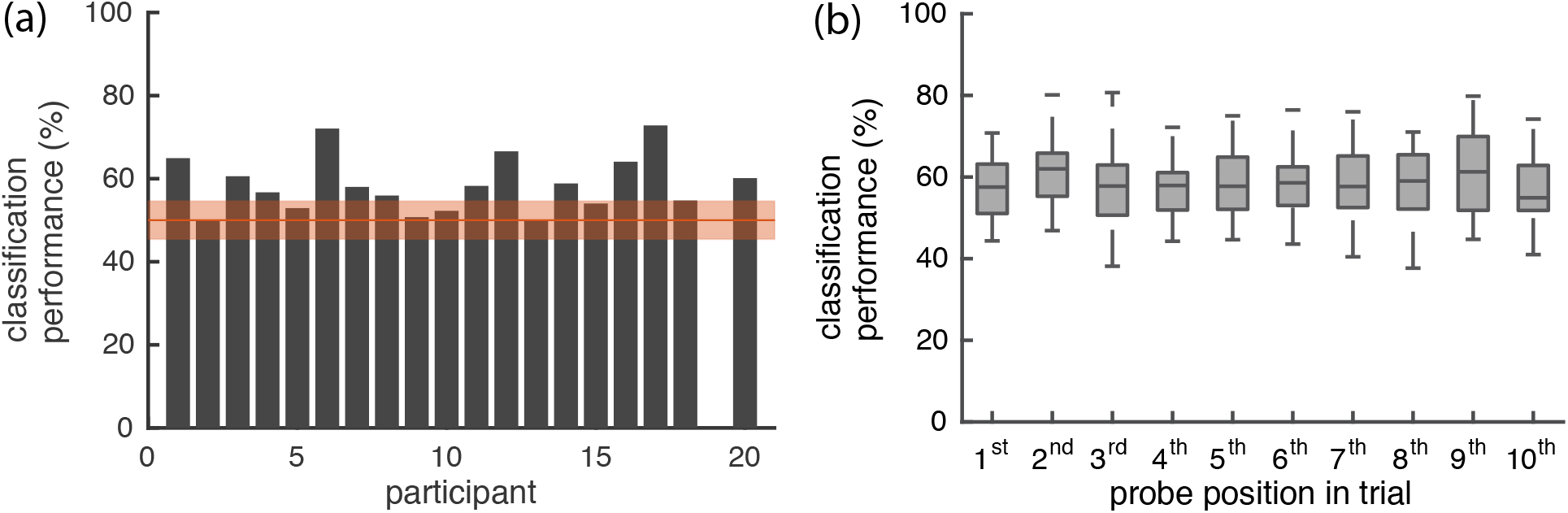
Classification accuracy of predicting (un)relatedness. (**a**) Classification accuracy per subject. Shaded in red, the 99.75% binomial confidence interval 50% accuracy (chance level; Bonferroni-corrected for number of subjects). (**b**) Classification accuracies for each position of a probe in the trial, across participants.

ERP plots for each individual subject are included in the Supplementary Information (Supplemental Figure S1), with their respective classification accuracy, as these may provide an idea of the variability across subjects in the ERPs themselves.

Using the behavioural measure, we can determine whether there is a relation between how well the participant’s behavioural responses agreed with our labels, and how well the relatedness of a probe can be predicted from their brain signals. The correlation between behavioural agreement and classifier accuracy, across participants, was moderately high (r = 0.58, p = 0.009, *α* = 0.05).

In Figure 3b we plot the classification accuracy based on where the probe occurred in a trial (i.e., the probe position). The boxplots show the distribution of performance across participants for a given probe position. There is no clear pattern visible between probe position and the accuracy of the prediction. To determine statistical (in)significance of any trend, we performed a permutation test, permuting the order of probe positions. Specifically, we compared the regression coefficient (of a line-fit) of the observed result against regression coefficients of results in which the order of probe positions was randomly permuted per subject (1000 permutations). The observed regression coefficient did not differ statistically from the coefficients from permuted results (p = 0.562; two-tailed alpha = 0.05).

### Time frequency analysis

To determine if there was any difference in oscillatory brain activity when contrasting the related or unrelated probes, we also analysed the data in the time-frequency domain. A cluster permutation test applied to the Time Frequency Representations (TFRs) of related and unrelated probes found a significant difference between the two conditions, identifying a significant negative cluster (p = 0.001, two-tailed, alpha = 0.05, n = 20)). To visualise this data, we subtract the TFR for related probes from the TFR for unrelated probes (i.e., unrelated – related) and plot this difference, averaged across electrodes, in Figure 4a. Clusters from a cluster-based permutation test are of limited use for determining which frequencies, timepoints, and electrodes in particular contribute to the significant difference, as the test does not control for the false alarm rate at this level (only the condition level contrast)^19^. Therefore, to aid interpretation, and in particular, to be able to compare our results to another study reporting effects in the time-frequency domain^17^, we performed two post-hoc significance tests, in which we isolated the alpha and beta frequency band respectively, and applied two cluster permutation tests, each on data averaged over the relevant frequency bands, and timepoints for the full trial (0 to 1.35 s). We Bonferroni corrected the significance values for performing these two tests, resulting in a significant effect for the beta band (p = 0.001), but with a non-significant cluster (p = 0.015) for the alpha band, at the two-tailed *α*/2=0.0125 level (two-tailed tests, Bonferonni-corrected *α* = 0.025, n = 20). These results are plotted in Figure 4b, with electrodes in a significant cluster marked with ‘ × ‘. In Figure 4a, we see an increase in power in the second half of the trial, that appears to be the strongest in the alpha band. However, the post-hoc significance tests determine that, though less pronounced, the difference in the beta band is, in fact, significantly different.

**Figure 4.**
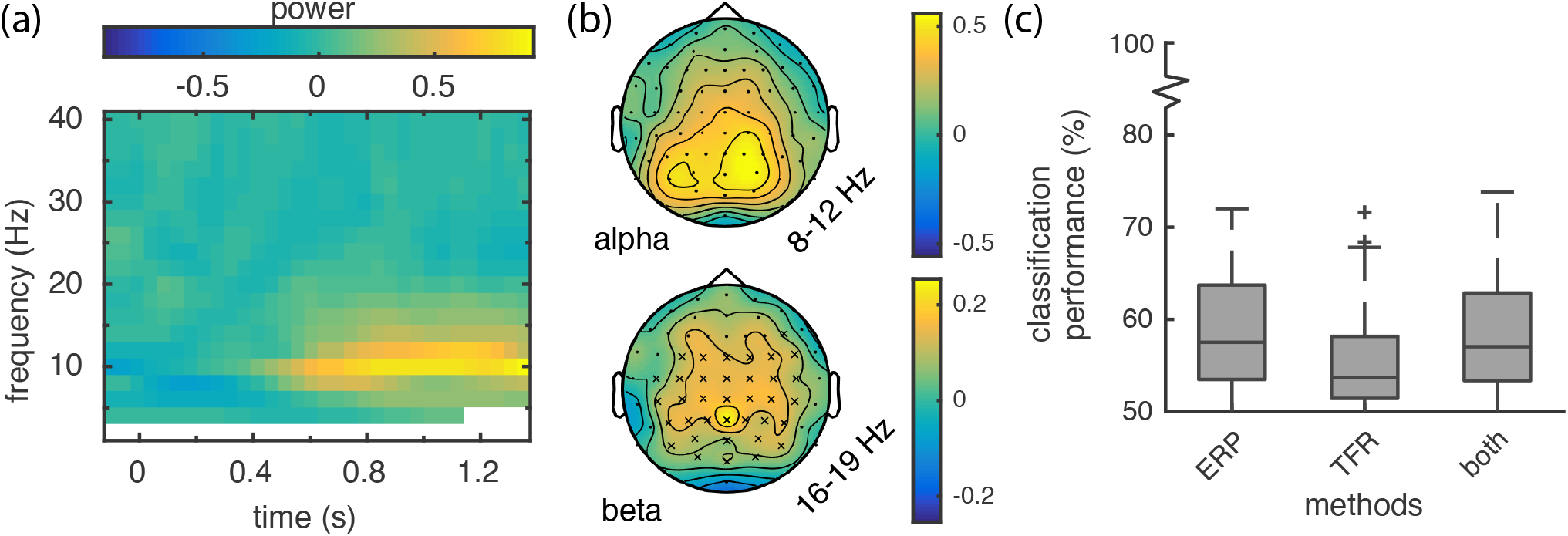
Results from the time-frequency analysis. (**a**) Grand average TFR of related probes subtracted from unrelated probes, averaged across electrodes. (**b**) Topographies for the alpha (8-12 Hz) and beta (16-19 Hz) bands. Electrodes belonging to an identified cluster are marked by ‘ × ‘. (**c**) Classification results of the ERP classifier, the TFR classifier, and a classifier trained on both feature sets.

To determine if this difference in oscillatory behaviour can be decoded from single probe presentations, we performed a classification analysis on this TFR data, analogously to the ERP data classification, and also created classifiers for both types of data combined. The results of this analysis can be found in Figure 4c. In the first boxplot we see a summary of the classification results shown earlier in Figure 3a, together with the TFR results and classifier accuracies from the combined data. It appears to be possible to predict the relatedness of a probe based on the time-frequency data for certain participants, but overall results look worse than for the ERP based classifiers. Furthermore, it looks like there is no benefit from including this data in addition to the ERP data when using a classifier.

### Accumulating predictions across probes in a trial

We have now generated predictions for single probes to estimate how well we can classify related from unrelated probes. However, for a BCI, information across multiple probes would need to be integrated to ultimately infer the original target. We can simulate such an analysis with our data, by using the consecutive predictions from a trial to predict which trial sequence they are from.

Each trial has a pattern of related and unrelated probes occurring in specific positions. We can try to predict from which trial a set of consecutive probe predictions was obtained, by comparing the similarity between the consecutive predictions and the true relatedness of all presented trials. That is, for each trial, we compute a similarity between the probe relatedness scores predicted by the classifier, and the ‘true’ relatedness expected for every possible trial, given the sequence of probe words. We then rank the trials based on this similarity (high to low). We can use this rank as an estimate of how well trials can be predicted from the sequence of probe predictions, by interpreting it as a percentile: a trial that receives a high percentile rank (e.g., 99th rank), obtained (among) the highest score on the basis of its predictions.

To show how this estimate changes as trials contain more probes, we start by only considering the first probe of each trial, then both the first and second probe, and so on, until all 10 probes are included in this analysis. In Figure 5, the results for the probe predictions on the individual participants’ data can be found, together with the results for randomly generated probe predictions (1000 randomisations). Note that the maximum percentile rank that can be obtained is dependent on the total number of possible patterns. This does not necessarily correspond to the total number of trials, as trials may have duplicate patterns. This ceiling is plotted in the figure, together with the number of unique patterns per number of consecutive probes considered.

**Figure 5.**
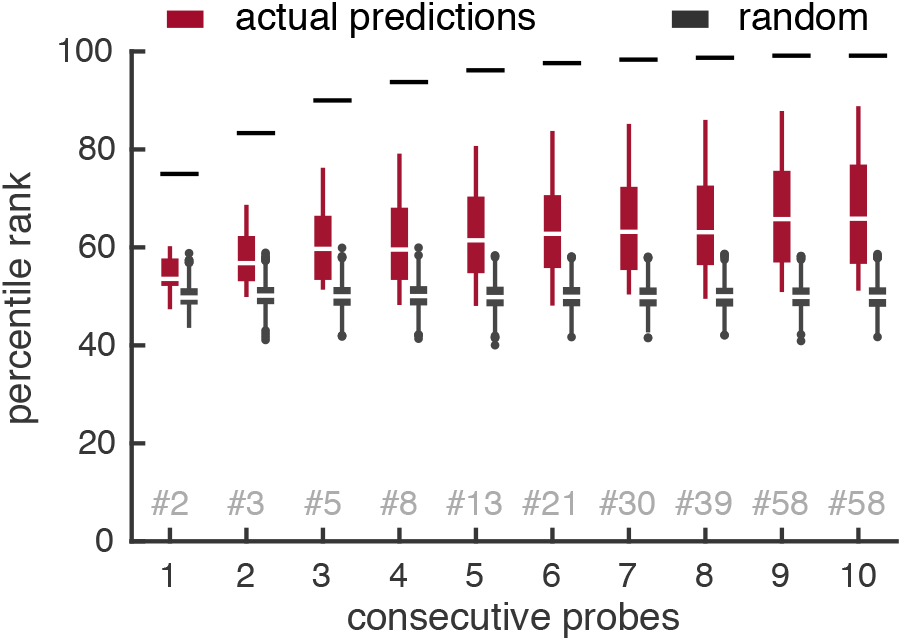
Percentile rank scores for predicting a trial, with varying number of consecutive probes considered. Percentile ranks based on either the participants’ probe predictions, or on randomly generated probe predictions are depicted. Horizontal lines denote the highest percentile rank that can be achieved given the number of unique trials. The amount of unique trials varies based on the number of consecutive probes that are considered, denoted in gray with a ‘#’.

The figure shows that for random predictions it does not matter how many predictions are accumulated for the percentile rank of the correct trial. For actual predictions, however, the (mean) percentile rank in which the correct trial is found increases when more consecutive probes are considered. There are also participants (in the tail of the boxplots) for which, even with 10 consecutive probes, the percentile rank cannot be distinguished from the random prediction data.

## Discussion

In this study we aimed to determine whether the N400 can be elicited reliably using multiple probes after an initial target. Overall, our results show that there is a difference in brain response to related versus unrelated probes, as evidenced by both the grand average ERPs and the classification results. Specifically, in the grand average ERP across probes from any position in the trial, we see an early component, more negative for unrelated stimuli, that matches the N400 in timing and location. Furthermore, the amplitude of this N400 is similar for probes appearing at the start and end of a trial (Figure 2), suggesting that the magnitude of the N400 does not reduce for probes occurring later in a sequence. In addition, classification results did not show any change with the position of the probe within a trial. Together, these results allow us to conclude that the N400 does not diminish over sequences of probes even when another, potentially distracting, semantic context was presented in between.

This result is consistent with a recent publication^20^, aiming to decode word relevance to a category of interest using sequential presentation of words and Fixation Related Potentials (i.e., ERPs time-locked to eye fixation of the stimulus). The authors found a more negative ERP for non-relevant words, similar to the N400. Interestingly, the results show markedly different brain responses for the online phase, where longer sequences were used (100 words), compared to the calibration phase (22 words). This may still be an indication that the N400 amplitude reduces for longer sequences (e.g., >10). However, with only a limited number of words used per category, stimuli were likely repeated frequently, which is known to cause a reduction in the N400 amplitude^21^.

An unexpected result in our initial analysis was an overlapping late-negative component occuring after related probes (see Figure 1a and Figure 2b). This could be a useful signal for BCI applications if it represents an additional signal to distinguish related from unrelated probes. Alternatively, as it appears most strongly for the final probes in a sequence, it may be an experimental artifact reflecting an expectation of trial end. In future work we aim to investigate these possibilities further. In this work, to alleviate the confound of this late component overlapping with ERPs from the subsequent stimulus, we post-hoc excluded unrelated probes that appeared after a related probe, reducing the number of unrelated probes in the final (53 probes) versus starting positions (104 probes).

Previous research has indicated that N400-tasks may also induce changes in power in the frequency domain^17^. When decomposing our data into the frequency domain, we visually observe *higher* power for related probes in the alpha and beta band from 600 to 1200 ms post stimulus presentation. However, only the beta band (16-19 Hz) increase was statistically significant. These bands were selected as in Wang et al.^17^ showed a *decrease* in beta power for in-congruent sentence endings (that elicit a more negative N400). This inconsistency in response suggests that either the role of beta power in relation to the N400 is different for sentence close-word paradigms, or that another aspect of our task (e.g., behavioural response preparation) results in these changes in beta power. These conflicting results, together with the classification analysis which showed no additional benefit from the inclusion of time-frequency data over ERP data only, suggest only limited usability of this response for a semantic BCI.

The main motivation for this research was to determine the feasibility of a sequential probing paradigm for a BCI that aims to infer the concept on a users mind through repeated presentations of probe words. In this respect, the fact that we did not find any indication of attenuation of the N400 is encouraging. Furthermore, while single trial classification accuracies are relatively low (50-72%), accumulating information across multiple probes demonstrably increases the information that can be inferred about the target (Figure 5). It is also clear that the ten probes are insufficient to identify which particular trial a set of probes belonged to. A semantic BCI will need to present more consecutive probes to obtain sufficient information. Furthermore, presenting random probe words will likely be inefficient. We believe that through intelligent selection of probes, significantly fewer probes will be required. For example, we can use information theory-based heuristics to maximise the information gained from a single probe by selecting probes such that the probe response will be most informative given the already acquired evidence (i.e., responses from previous probes).

For approximately a third of the subjects, the relatedness status of a probe-prime pair could not be predicted from the EEG with better than chance accuracy. For comparison, in the prime-probe study by Geuze et al.^16^, all participants’ classifications results exceed their chance threshold (p < 0.05). However, when correcting their binomial confidence interval for multiple comparisons, as we did here, their chance level would be at 57% with only 8 (67%) participants exceeding this threshold. It appears that the N400 cannot be detected reliably in everyone, which is not an uncommon problem for BCIs (see e.g.,^22^ for a discussion). Furthermore, Cruse et al.^23^ have demonstrated that detectability of the N400 in individuals depends on the experimental parameters, finding that both the task: behavioural, mental, or passive and the type of relatedness (semantically similar vs associative), impacts the ability to identify the N400 in individual subjects. This sensitivity to task parameters suggests that this is in part a signal-to-noise problem. Furthermore, the correlation (r = 0.58) we found between the behavioural agreement with the labels and the N400 classifiability, additionally supports the notion that the degree of attention to the stimulus is important.

In the end, while the current study presents a strong case for the *theoretical* feasibility of a BCI that infers the concept on a user’s mind, the *practical* applicability of any application will depend largely on the time required to (accurately) select a concept, and the size and type of vocabulary that concepts can be selected from. The time-to-selection will depend primarily on the number of probes required to achieve the desired accuracy, which is difficult to estimate. For instance, 50 probes, presented at the current SOA (1350 ms) would require a little over a minute for selection, which may be acceptable for certain applications. With regard to vocabulary, this approach is inherently geared towards selecting words (i.e., concepts) that have semantic content, such as nouns, verbs, adjectives, etc. This potentially excludes the selection of function words (e.g., conjunctions, prepositions and pronouns), precluding the ability to construct sentences to some degree.

To what degree this affects practical applicability may depend on the target application. For instance, the proposed BCI could be used to restore the ability to communicate to specific patient groups. While it is unlikely that a semantic BCI will reach the performance of the fastest—but gaze-dependent—speller BCIs, by using spoken words they may provide a viable alternative to gaze-independent spellers. These achieve lower communication speeds, but offer a method of communication to patients that can no longer direct their gaze. While not tested here, the N400 has been elicited reliably in the auditory domain^12,13^, suggesting that auditory probes may be used as an alternative to textual stimuli. Nevertheless, sentence construction will prove difficult if function words cannot be selected, though a semantic BCI may still serve as an additional tool that can be used in conjunction with a speller system, as an important advantage of this probing paradigm is that it only requires the user to concentrate on the to-be-communicated concept, rather than composing a word character by character. A further advantage of the use of auditory stimuli in this paradigm is that it does not require the user to be literate. In the same vein, picture stimuli could be used for illiterate users, as these can also be used to elicit the N400^24^.

Alternatively, semantic BCIs may be able to offer assistance to users with anomia (an inability to name words)^8^. This BCI may provide an intuitive way for them to retrieve their intended word. However, it remains a question whether patients with anomia will exhibit a normal N400. For instance, in patients with aphasia, whom frequently deal with anomia, the N400 amplitude can be reduced, the degree of which appears linked to their comprehension ability^25,26^.

In summary, our results show that we can decode responses of relatedness from EEG, and we find no indication that this response attenuates across multiple consecutive probes. Without evidence of attenuation, and with an illustration of how evidence from multiple consecutive probes can be accumulated to identify an initial target, we see a clear path toward a semantic BCI that uses repeated probing to infer the concept on a user’s mind.

## Methods

### Participants

Twenty-one participants took part in the experiment (12 female, 9 male), ranging in age from 18 to 61 years old (mean age 29). Participants were only eligible to participate if they were native Dutch speakers and reported to have no reading problems (e.g., dyslexia)

One participant dropped out of the experiment due to excessive sleepiness. Her data are not included in the analysis. Response data are missing for participant 9, block 6, due to a technical problem, and there is no data for participant 19, block 6 (due to initial cap fitting difficulties, the experiment could not be completed in the allotted time-frame).

All participants provided informed consent prior to participant, and the study was approved by and conducted in accordance with the guidelines of the Ethical Committee of the Faculty of Social Sciences at the Radboud University.

### Task

In the experiment participants completed 212 trials, each consisting of a target word, followed by one to ten probe words. For each trial, participants were tasked to remember the target word, and subsequently mentally assess for each probe whether or not it was semantically related to the target. At the end of a trial participants were prompted to specify by button-press the relatedness status of the most recent probe (left = related, right = unrelated). 50% of trials contained ten probes, while the remaining 50% contained an equal distribution of one to ten probes. Trials with fewer probes were included to ensure that participants would mentally evaluate all probes, and not only those for which they could predict that a question would follow.

Participants were instructed to respond as fast as possible when prompted with a question, and received their reaction time as feedback (this feedback was coloured using a gradient: green to white to red at 700 ms, 950 ms, 1200ms, respectively). This feedback did not reflect ‘correctness’ of the choice as relatedness judgements can vary between individuals. To increase salience and facilitate memorisation, participants were asked to pronounce the target word during its presentation.

Each target word was presented for 2000 ms, followed by a 1000 ms interstimulus interval (ISI). Each subsequent probe was presented for 350-370 ms (jittered duration) and also followed by a 1000 ms ISI. The question prompt was displayed after the last ISI, and was replaced by feedback on buttonpress. Across the 212 trials a total of 539 related, and 1072 unrelated probes were presented (33.5% related).

In our analyses we compare probes occurring in the first position with probes occurring in the ‘final’ position (i.e., 9^*th*^ and 10^*th*^ position). Note that we define the ‘final’ category to consist of two probe positions, in order to get an equal number of brain responses, as only 50% of trials consisted of the full 10 probes. With this analysis in mind We designed trials such that there was an equal distribution of related and unrelated probes for the first and final position(s). Finally, these trials were set up such that no two related probes were ever presented in a row, as a brain response signifying relatedness in such a case could be purely from the pairing with the previous stimulus. The percentage of related probes per probe position can be found in Figure 6a.

**Figure 6.**
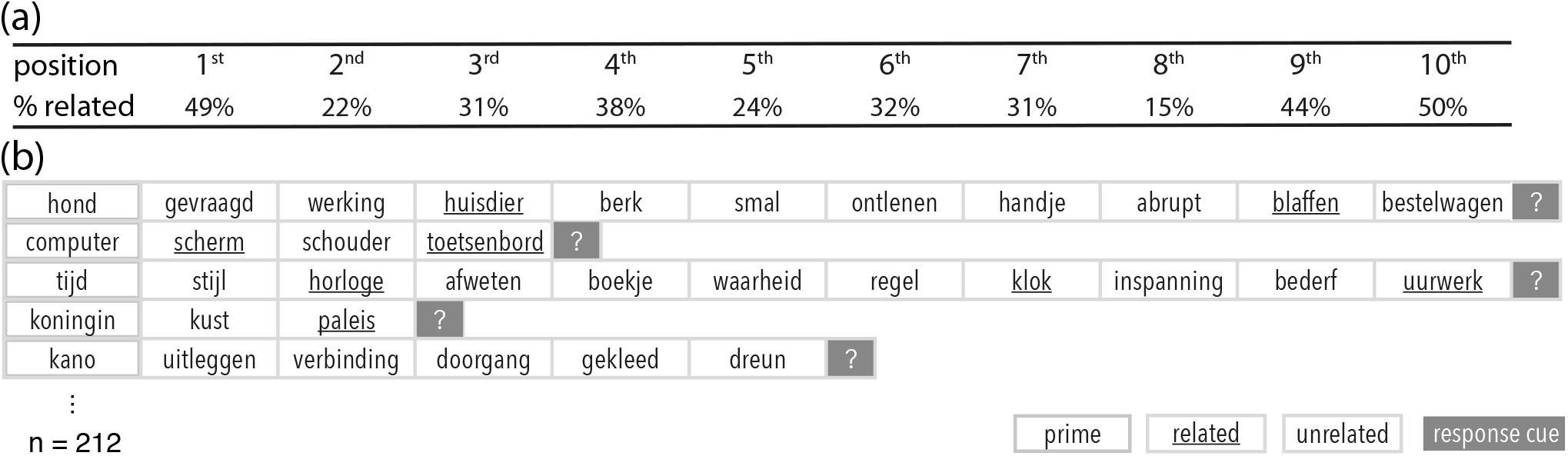
Trial design with example stimuli. (**a**) Percentage of related stimuli per probe position in the trial. (**b**) Example trials: A trial consists of an initial target, followed by up to 10 probes. A probe is either related or unrelated to the target. Each trial ends with a behavioural prompt.

Stimulus words were obtained from the Leuven association dataset^27^. This Dutch dataset consists of cue-words and responses from people who were asked to specify (up to) three words that a given cue-word brought to mind. In our design we used Leuven cue-words as targets, and Leuven response-words as (related) probes. We then used the CELEX Dutch Wordform database^28^ to obtain words unrelated the the targets, to use as unrelated probes. Probes never occurred more than once across all trials, as repetition of concepts has been shown to affect the N400 amplitude^21^, though some targets also occurred as probes. Unrelated words were selected to approximate the related probes in distribution of both word length and word frequency. Word frequency in particular, is known to affect N400 amplitudes, with words that occur frequently in a language eliciting smaller N400s. Across all probes, the mean (log) word-frequency was 1.50 (sd = 0.65) for related, and 1.49 (sd = 0.63) for unrelated probes. On average the length of a related probe was 5.68 characters (sd = 1.98), and an unrelated probe 6.13 characters (sd = 1.92).

Stimuli for a few example trials are depicted in Figure 6b. A detailed description of the selection procedure and a full list of stimuli can be found in the Supplementary Information.

Trial order was randomised per subject.

### Equipment

EEG was recorded with 64 sintered Ag/AgCl active electrodes (BioSemi, Amsterdam, The Netherlands), placed according to the international 10-20 system, at a sampling rate of 512 Hz. The Biosemi ActiveTwo system uses a Common Mode Sense (CMS) and a Driven Right Leg (DRL) electrode, instead of a ground electrode. The recorded signals reflect the voltages between each electrode and the CMS, which can subsequently be re-referenced to any electrode(s) of choice. We placed two electrodes on the mastoids for this purpose and used four further electrodes to measure vertical and horizontal EOG.

### Data Analysis

#### Preprocessing

Data were recorded in 6 blocks of 35 trial sequences each. These data blocks were loaded, highpass filtered at 0.1 Hz (4th order Butterworth filter)^29^, and then sliced into segments, from 1.5 s prior to and 2 s after each probe presentation. Then, data from different blocks were combined and downsampled to 256Hz. All electrodes were de-meaned and re-referenced to the average of the mastoid electrodes. The four EOG channels were then regressed-out of the EEG channels to remove any influence from eye movements or blinks^30^. The EEG channels were subsequently lowpass filtered at 40Hz.

After these preprocessing steps, any trials or channels with abnormal activity were marked for removal (trials) or interpolation (channels). A trial or channel was considered abnormal when the variance of the given unit diverged more than 3.5 standard deviations from the median of all units (channels or trials). This was repeated, excluding previously marked units in the next iteration, until no units were considered abnormal (max 6 iterations). This resulted in between 12 and 156 trials and up to 4 channels to be marked per participant. Any identified bad channels were replaced with an interpolated ‘virtual channel’ using a spherical spline interpolation^31^, while bad trials were marked to be ignored during grand average calculation and training of classifiers. Finally, trials were baselined to a period from −100 ms to 0 ms from stimulus onset.

#### Grand Averages

For brain responses in the time domain, we obtained grand average ERPs by averaging across relevant probes, and then averaging across subjects. The data for the grand average ERPs were low-pass filtered at 20Hz, prior to averaging, for plotting purposes (smoothing). To analyse brain responses in the frequency domain, the data from each trial was decomposed into frequency bins of 2 Hz, from 2 to 40 Hz. The power in these bins was calculated for each 50 ms, starting from 0.5 s prior to probe onset to 1.35 s after, using a Hanning window and a frequency dependent window length (ranging 3.5 s - to 175 ms respectively). These Time Frequency Representations (TRFs) were then averaged across participants to obtain grand average TRFs.

#### Significance testing

To determine whether brain responses between conditions (e.g., related vs unrelated) were significantly different, we used non-parametric cluster-based permutation tests^18^ (as implemented in Fieldtrip^32^; www.ru.nl/neuroimaging/fieldtrip). Such a test allows for the combination of information across electrodes and timepoints, to increase sensitivity of the statistical test, without having to correct for multiple comparisons with respect to those aspects. These tests were performed using a within-subject design, and a dependent samples t-test as test-statistic. Note that this non-parametric cluster-based permutation test does not rely on assumptions with regard to the distribution of the data, regardless of the chosen test-statistic (the test-statistic is merely used to quantify the difference between datapoints). In all cases, we performed a two-tailed test, (tail-corrected alpha = 0.025), using 1000 permutations, supplying all channels, and timepoints and/or frequency bins as specified.

#### Classification

For classification analyses on data in the time domain, the preprocessed data were further downsampled to 64Hz and low-pass filtered to 15Hz to reduce the amount and complexity of data passed to the classifier and prevent overfitting. We use a classification pipeline identified as robust for different types of ERPs^33^, consisting of a spectral filter, a spatial whitening and classification using a regularized classifier. Specifically, the preprocessed data from time 0 to 1s and all remaining trials and channels (already spectrally filtered), were spatially whitened, and subsequently classified using a L2-regularized logistic regression. The regularization parameter was optimized using a 5-fold (nested) crossvalidation. For classification analysis on data in the time-frequency domain, preprocessed data were again decomposed into frequency bins of 2 Hz (2-40Hz), but here the bins were calculated for each 100 ms from probe onset (0 s), until the end of the trial (1.35 s), using a Hanning window and a frequency dependent window length (1 s - 175 ms respectively). These time-frequency data were then processed using the same classification pipeline.

To estimate classification performance, the data were separated into crossvalidation folds of training and test sets, where all but one trial sequences were used as training data, and the data from probes in the excluded trial sequence were used as the test set. Only trial sequences in which ten probes were presented were used as test sets, to avoid biasing the results toward responses to early probes. Trials that were marked for removal during preprocessing were excluded when occurring in a training set, but were included when part of the test set (in an online BCI application poor data quality does not always preclude an accurate prediction).

## Supporting information

## Acknowledgements

We would like to thank James McQueen for valuable feedback on an earlier draft of this paper.

