## Supplementary material for "Semantic Probing: Feasibility of using sequential probes to decode what is on a user’s mind"

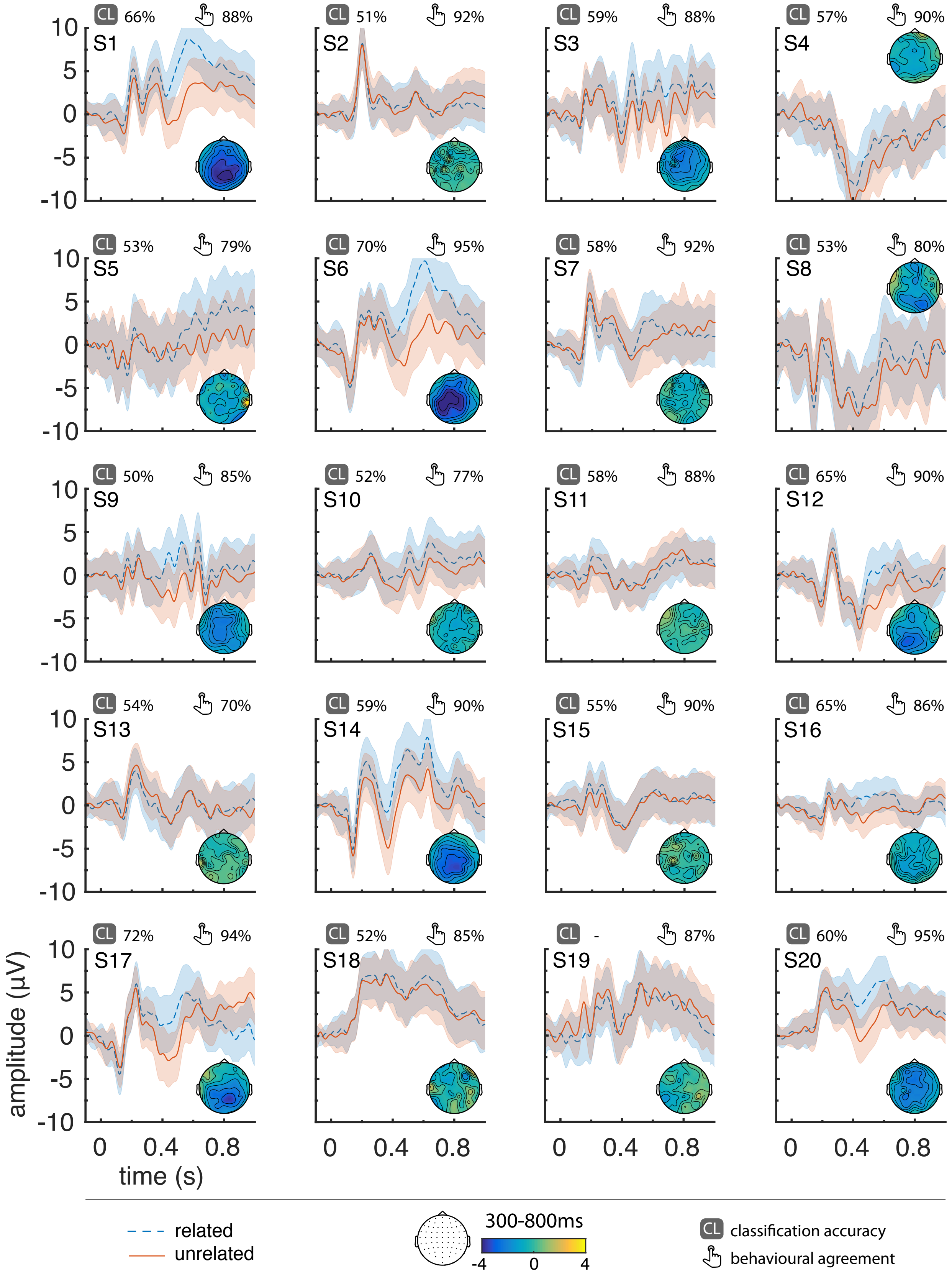


**Supplemental Figure S 1. Individual ERPs with shaded standard deviations.** Spatial topographies from 300-800ms are included. Classification rates and behavioural agreement with the ground truth labels are specified for each subject.

### Supplementary Methods: Description of stimuli

Stimulus words were obtained from the Leuven association dataset (Deyne and Storms; 2008), and the CELEX dutch wordform database (Baayen et al.; 1995; celex.mpi.nl). The Leuven dataset is a Dutch word-association database that consists of cue-words and responses from people who were asked to specify (up to) three words that a given cue-word brought to mind. The database is a raw collection of the human responses, and can contain misspelled words or nonsensical responses. To construct a sensible stimulus, set we filtered the responses in this database to contain only headwords (i.e., how a word occurs in the dictionary), by matching it against a CELEX list of words with the 'headword' property.

The Leuven cue-words could then be used as targets, and the responses to these cue-words as related probes. To determine how targets and probes could best be allocated to trials, we constructed a trial framework such that 50% of trials contained ten probes, and the remaining trials contained an equal distribution of one to ten probes. We ensured that the first and final ($9^{th}$ and $10^{th}$) positions contained an equal number of related and unrelated probes. The remaining probe positions (2nd through 8th) were designated to contain a related probe with 1/3 probability, while ensuring no two related probes occured consecutively.

Having now specified the number of related and unrelated probes required for a given trial, we could now assign targets (i.e. Leuven cue-words) with a sufficient number of related probes (i.e., Leuven responses) to each trial. To avoid repetition effects, probes that were used in one trial, did not occur in any other trial as a probe. However, some probes did also occur as a target. In such cases, the trial in which it occurred as a probe would be presented earlier in the experiment than its occurrence as a target, where possible. Unrelated probes were drawn from the previously mentioned CELEX head-word list, but now with the already selected related probes and targets removed. For each trial, we made sure that the selected unrelated probes had not been given as a response to the cue that corresponded to that trial's target, according to the Leuven database.

Furthermore, to make sure the unrelated probes had similar word length and word frequency, we first sampled a subset of CELEX words to match the distribution of word lengths to that of the already selected related probes. Then, when assigning unrelated probes, these probes were sampled to match the distribution of word frequencies of the related probes. Across all probes, the mean (log) word frequency was 1.50 (sd = 0.65) for related, and 1.49 (sd = 0.63) for unrelated probes. This resulted in average word lengths of 5.68 (sd = 1.98) for related, and 6.13 (sd = 1.92) for unrelated probes.

Finally, the resulting trials were manually checked to remove any issues (e.g., probes assigned as unrelated being related by chance).

The trials were then shuffled per participant, in such a way that probes that were also targets, first appeared as a probe before they did as a target. Note: due to some three-way connections across trials this was not *always* possible.

### Supplementary Methods: List of stimuli

Below, a complete list of stimuli presented in the experiment. Targets are **bolded**, and related probes are underlined.

**slapen** - schoot - dromen - edel - nacht
**wandelen** - koel - chip - park - zuiver
**verpleegster** - wand
**hoofd** - denken - omgeven - hersenen - grootte - engel
**zilver** - juweel - vijand - vertrek - ketting - heuvel - body - genegenheid - druppel
**leger** - soldaat - strak - belonen - tank - ark - geweer - weten
**molen** - graan - aurora - wind - bekken - meel
**banaan** - aap - emmer - zes - krom
**sport** - tennis - apart - openen - voetbal - aangifte - zwemmen - aanmoedigen - zweet - buik - lopen
**snijden** - bodem - scherp - mate - mes - volk - bloed - aankomen - cijfer - atelier - vlees
**vinger** - bouwer - weigeren - erf - duim - behang - gedrukt - winst - dank - huidig - wijzen
**berg** - top - steun - bui - natuur - flits - buit
**wandelstok** - basis - zijde - jongere - wandelen - badkuip - opa - aula - heten - bergen - bonken
**gordel** - veiligheid
**revolver** - charme - bot - roman - cowboy - grijns - staf - droevig - zeven - bestuur - moord
**handtas** - wilde - rommel - stem - leer - beest - geld - dertig - veranderen - vrouw - belofte
**hamer** - kloppen - boeg - getuigenis - gewijd - pogen - ruim - komen - gade - spijker - verenigen
**keuken** - tafel
**toetsen** - leren - afkeuring - bezocht - verhouding - stress - begeerte - veel - computer - bel - examen
**bar** - drinken - meemaken - tap - bepalen - kruk - zingen - behoefte - gedaan
**zitten** - staan - poging - consul - bank - dunne - vlucht - totaal
**sigaret** - roken - belijden - ongezond - armoedig - kanker - eindeloos - elleboog - beschouwen - tabak - sluiten
**gewicht** - kilo - durven - symbool - brein - weegschaal - bannen - jagen - juli - barkeeper - zwaar

**aarde** - simpel - beverig - bakje - bol - bevoegde - slag - planeet - lijn - wereld - keren
**dierenarts** - erkenning - stuur - knaap
**blazen** - beklag - zakken - handeling - ballon - gids - conceptie
**koningin** - kust - paleis
**boek** - bibliotheek - afweer - ontspanning - achterste - spannend - baron - roman - gebied - waar
**hoorn** - driftig - telefoon - lijf - blazen - evenwicht
**boter** - smeren - autobus - vet - autonomie - bakken - avenue
**touw** - eeuwigheid - koord - anus - trekken - jaar - arend - knoop - leeg - klimmen - boord
**naaimachine** - naald - conditie - stikken - betoging - wet - emotie - automaat - banning - garen - boerin
**vuil** - modder - blos - afval
**vrouw** - genoemd - mooi - afwegen - man - afbeelding - moeder - geleid - afnemen - erg - kind
**pruim** - integratie - lichaam - gang - vrucht - actief - paars - bazin - barst - pit - beslissend
**stofzuiger** - gezet - vuil - arena - armoe - stof - rijke - lawaai - binden - golf - tapijt
**beeld** - noemen - schaal - plaat - losmaken - kunst - beheerder - foto - meest - standbeeld - juist
**bijl** - brandstof - dulden - opvallen - proeven - buisje - bindmiddel - binnenste - lijk - dozijn - hakken
**bliksem** - los - assistentie - regen - bijbels - donder - relatief - hut - brullen - onweer - adem
**oud** - afbreken - afmeting - grootouders - afgrond - beleid - jong - adoptie - oma - anoniem - versleten
**fruit** - peer
**pop** - gezegd - speelgoed - aangeven - meisje - bezinning - spelen
**kano** - uitleggen - verbinding - doorgang - gekleed - dreun
**oester** - schelp - attent - parel - activa - akker - slijm - vorig - paniek - bezoek - zout
**kleur** - zonde - ruzie - hees
**fabriek** - werken - genieten - werk - abstractie - rook - jongeman - arbeider
**boom** - tak - gering - gereed - letten - stam - westen
**cowboy** - bijlage - zaal - hoed - kil - oktober - neiging
**tand** - balsem - speciaal - broer - leveren - concern - tandarts - opvatting - gebit - verlenen
**proper** - net - bestand - bevalling - schoon - leven - bizar - bloeddruk - diplomaat - netjes - dienen
**rechter** - toga - eigen - gevel - ban - advocaat - reeks - rechtbank - gerust - belast
**peer** - appel - voorzien - brons
**staart** - bidden - paard - haat - hond - atmosfeer - kat - amulet - leerling - luid - lang
**panda** - karakter - kern - beer - informatie - kwartier - bamboe - gaande - boa - deftig
**handdoek** - doorbreken - normaal - drogen - zaak - afwas - lieden - revolutie - beding - droog - diepte
**pasta** - hoop - knie - kosten - schok - wagen - spaghetti - beleven - burger - stoten - saus
**wol** - brandend - schaap - ballet - onverwacht - relatie - trui - maal - pakken - breien - bits
**reis** - braken - drager - vak - hertog - verstand - voldoen - fabriek - klacht - ver - firma
**kleerkast** - kleren - afwijking - aandoen - slaapkamer - bemanning - achterhoede - kledij - meneer - moment - kapstok
**dief** - bezield - vlag - oprit - cacao - overgaan - vermogen - storten - bijval - camping - stelen
**hond** - gevraagd - werking - huisdier - berk - smal - ontlenen - handje - abrupt - blaffen - bestelwagen
**inkt** - vlek - bokser - trek - luiden - mand - vulpen - werker
**munt** - thee - buffer - mus
**lucht** - vliegtuig - adres - ademen - lichten - kaal - min - berouw - laten - wolk - twintig
**boerderij** - boer - geestelijke - koe - baken - varken
**teen** - poos - stimuleren - oever - radio - vrede - gunst - wijd - boze - voet - bushalte
**parachute** - rollen - ego - spannen - inclusief - donkergroen - ruw - spiegel - continu - springen - streven
**clown** - grappig - bitter - gauw - circus - compleet - leunen - lachen - ingaan
**exotisch** - cocktail - ader - zon - bedragen - ananas - dichtbij - eiland - aanpak - barsten
**tas** - grap - buikpijn - zak - visie - wonder - puur - bewonderenswaardig
**computer** - scherm - schouder - toetsenbord
**noordpool** - zuidpool - fort - rapport
**zien** - kijken - kraken
**punt** - komma - haven - potlood - belang - zin - geest - axioma - dekken - bewijzen - asiel
**scheermes** - schuim - klasse - bukken - baard - balie - snijden - artiest - doos - benoemen - gebogen
**akker** - beschut - veld - aroma - eis - kruipen - land - gemiddeld - doen - dapper - landbouw
**hand** - vinger - wil - norm - genoegen - beraad
**garnaal** - zee - aanzienlijk - functie - lekker - aanleren - tomaat - baseren - vis - ruimte - grijs

**slim** - intelligent - weggaan - behendig - dom - gemak
**zwaan** - uiting - blok - kom - zenden
**bezem** - heks - bil - bedaren - omvang - partij - benzine - steel - datje - vaak - vegen
**wesp** - steek - liegen - bedrog - bij - aanbod - steken - asbak - orde - anjer - angel
**tram** - anima - sporen - duif - atleet - stad - kist - behoren - spoor - krant
**televisie** - nieuws - dubbel - film - tamelijk - programma - exact
**wiel** - kar - benul - voornaam - band - bondig
**rat** - vies - genomen - riool - fris - staart - doden - eeuwig - hoeven - muis - bewegen
**meloen** - oranje - type - dorp - zomer - balans - sappig - bouw - leider - rond - boeien
**drinken** - dorst - citeren - cola - achtergrond - beroep - lenen - acceptatie - alcohol - verder - glas
**raam** - bon - inzien - twijfel - venster - bronzen - commando - vullen - boosheid - proef - buiten
**wild** - hert - moed
**hek** - afsluiting - verte - baret - omheining - niveau - liggend
**bril** - westers - volwassen - zuster - lezen - fijn - bewoner - montuur - long - boel - glazen
**kooi** - gevangen - dragen - museum - week - leeuw - loop - gevangenis
**muur** - hoog - bestaan - ambitie - cement - alarm - affaire - afzet - baksteen - balk - hard
**jeuk** - krabben - mens - uitslag - ideologie - bast - elf - beleg - mug
**post** - postbode - arsenaal - brievenbus - dode - zwarte - bejaard - aspirine - postzegel - degelijk
**tuin** - behoud - planten - gezag - gave
**priester** - bouillon - tevreden - bruto - bestuursrecht - kruis - voedsel
**lade** - kast
**kampvuur** - warmte - druif - gezelligheid - drie - gezellig
**handschoen** - koud - contact - warm - heer - winter - agent - sneeuw - krijgen - hand - afkeer
**veel** - bankje - weinig - huren - massa - barman - meer
**taart** - ergeren - boef - slagroom - ford - aanpassen - keel - verjaardag
**station** - trein - democratie - beoordeling - wachten - rol - perron - formule
**oor** - vinden - later - horen - geval - luisteren - baai - affectie - muziek - een - geluid
**knol** - wortel - pagina - jeugd - gezocht - grond - dek - sprake - fors - aardappel - lot
**baas** - vroeger - streng - leggen - chef - boog - beurs - bureau - orgaan
**verkeer** - facet - file - kopje - ziekte - duidelijk - bezwaar - drukte - ding - vereniging - druk
**geweer** - schieten - afgemeten - middel - kogel - daad - as - jacht - aanplant - bond - dood
**koe** - weide - leraar - akte - gras - context - kalf - beheer - bestuurlijk - wei - sprong
**duivel** - slecht - bijten - bloedig - hangen - blank - dun - hel - vast - satan - vorm
**schild** - positie - bereik - deugd - ridder - baarmoeder - cent - pad - diep - bescherming - stand
**spijker** - kop - baas
**kanarie** - deeg - vuist - kooi - verstaan - zuiden - fluiten - boeten - border - bod
**glijden** - slee - juni
**koning** - koningin - eindje - dominee - kroon - rustig - troon - alfa
**broek** - rok - ouder - noodzaak - strijd - tien - grote - rits - sterk - jeans - redden
**blok** - billijk - collectie - bindend - beton - vormen - bisdom - studeren - bacon - breedte - been
**uniform** - leger - aanstelling - afkomst - politie - half - school - aanvraag - zit - blauw - auteur
**peper** - kwaad - gedaante - pikant - denkend - titel
**bok** - jasje
**vulkaan** - lava - azijn - gaan - bede - huid - uitbarsting - doel - gast - berg - negeren
**schoorsteen** - gezin - toon - zwart - formeel - isoleren - sinterklaas - oude - kijker - roet - begrip
**tang** - duiden - botsing - ijzer - goed - aquarium - hoek - gereedschap - beestje - werktuig - best
**robot** - metaal - zak - atoombom - eigenschap - toekomst - gebruik - machine - begerig
**engel** - hemel - cursus - arrestatie - lief - ambt - duivel - madame - duren - god - bladzijde
**das** - feest - bak - hemd - taak - dal - leiden - kostuum - kader - pak - oog
**trouwen** - ring - alfabet - antenne - kerk - eng - dagboek - liefde - begaan - jurk - arrogant
**kapitein** - gebrek - vijfde - schip - tocht - stevig - noorden - vaag - bekleed - matroos - rij
**badkamer** - douche - grens - wassen - vier - bad - model - aanzien - afslag - handdoek - ervaren
**helm** - graad - brommer - expert - motor - bijstand - collega - oor - blom - veilig - externe
**ontbijt** - meten - boterham - assistent - koffie - bewind - brood - bedrijvigheid - kennis
**pad** - groei - kikker - sturen - glibberig - astma - andijvie - blik - vijver - angina

**vonnis** - gerecht - verband - rechter - beitel - beklemming - blij - merken - nerveus - straf - bakkerij
**bom** - explosie - afhouden - beiden - wensen - oorlog - bende - ontploffing - breed - binnenlands - geweld
**yoghurt** - wit - officier - melk - afhalen - fruit - voelen - suiker - avontuur - aankoop - gezond
**eiland** - vrezen - dimensie - onbewoond - kleden
**eiwit** - kip - bootsman - eigeel
**beha** - kant - helder - ondergoed - berber - klagen - barok
**knopen** - opzicht - touw - zetten - jas - debat
**knippen** - schaar - praten - enorm - papier - enig - hanteren - kapper - dalen - haar - afname
**pot** - deksel - kiezen - ketel - fase - betoog - agenda - aarde - berekenen - afdronk - plant
**gitaar** - buffet - defensie - interessant - schuin - snaar - opmerking - prins
**chips** - broche - slepen - paprika - cilinder
**opstaan** - intiem - inwoner - elite - vroeg - den - bosrand - eens - kring - wekker - domein
**rad** - fel - wiel - barbaars - heet - kermis - koffer - zoon - draaien - klas - fortuin
**tafel** - tafelkleed - verwijderen - rede - nodig - poot - binnenland - nek
**omelet** - ei - alibi - grof - geel - allergisch - pan - dag - ogenblik - spek - abdij
**dieet** - beige - stijgen - experiment - beloven - citroen - honger - duwen - bedolven - mager - nota
**papegaai** - boeddha
**aas** - gier - benadering - worm - aanwending - vissen - droom - absurd - kaarten - aapje - hengel
**schakelaar** - geloven - lamp - baden - brengen - chaos - knop - beeld - aan - atoom - elektriciteit
**ruiken** - geur - grondslag - parfum - aangeschoten - huwelijk - stank - missen - eten - afwezig - bloem
**divan** - begeven - liggen - anker - zetel - verlies - rusten - brutaal - ambassade - reden - slapen
**hak** - beetje - winkel - erkend - bijl - verzet - beha - heffen - kwestie - hiel - vergelijken
**weg** - baan - schaduw - genie - broekspijp - bouquet - dialect - worden - mama - bazaar - straat
**magneet** - nood - wenden - aantrekkingskracht - hoesten
**anker** - oosters - doolhof
**vuist** - circuit - kapot - prof - maan - bestuurslid - naakt - vechten - cyclus - mei - verslag
**trap** - gebed
**kraam** - nadruk - gretig - waarneming - koers - provincie - riks - pastoor - blaas - markt - buis
**kogel** - rijzen - pistool
**naaien** - klaar - naaimachine - stap - kleding
**ladder** - inhoud - geniaal - dak - champagne - boezem - bas - pop - gigantisch - trede - onzeker
**microscoop** - onderzoek - achttiende - acceptabel - arriveren - dame - klein - gat - wetenschap - stukje - biologie
**venkel** - crisis - kruid - zuchten - betaald - soep - gevoel - groen - aartsbisschop - groente - gevaar
**dessert** - taart - fantastisch - bedachtzaam - chocolade - factor - beide - zoet - bloei - pudding - naam
**instrument** - hoge - gitaar - dicht - viool - glanzen - trompet - globaal - kamer - bereikbaar - piano
**neus** - animo - gezicht - beter - ruiken - behouden - glimlach - zakdoek - beoordelen - beroemd - groot
**ridder** - beugel - doctor - harnas - casino - doorn - zwaard - moeten - blaadje - flink - prinses
**ketting** - blinde - zaag - breuk - hals - keer - pers - benoeming - ernst - rode - goud
**schaken** - bochel - moeilijk - cabine - plicht - joods - koning - oorzaak - vol - spel - stroom
**tijd** - stijl - horloge - afweten - boekje - waarheid - regel - klok - inspanning - bederf - uurwerk
**bikini** - zwembad - bevriend
**tijger** - geografisch - wild - typisch - geheel - uitgaan - gevaarlijk - badwater - begroeid - blouse - jungle
**kussen** - capitulatie
**ondergronds** - metro - afgunstig - beker - kelder - beroven - derde - reis - donker - bruikbaar - tunnel
**sleutel** - consumptie - wens - slot
**lepel** - apathie - vork - maand - bord - papa - bestek
**speer** - falen - actie - werpen - bewering - punt - raad - hebben - opgeven - wapen - verhaal
**palet** - hout - bestaand - analoog - schilderen - allereerst - verf - hulp - aardgas - schilder - album
**veer** - baren - zacht - dof - neger - pluim - banaal - vogel - argeloos - nadenken - schrijven
**woestijn** - kameel - beuk
**wolf** - janken
**glibberig** - alomvattend - vallen - enkel - ijs - angst - deel - glad - aardig - anarchie - slang
**kluis** - graaf - code - vloer - wang
**bakken** - held - tonen - bezetter - eisen - cake - prijs - oven - beul - bakker - heftig
**alcohol** - kennen - bier - behulp - wijn - afwijzing - drank - matig - dronken

**bromfiets** - front - jack - damp - vochtig
**snavel** - vangen - datum - eend - hek - helling - dij - dichter - geloof - terras - bek
**chauffeur** - mogen - auto - beginnen - bus - katholiek - afgewogen - taxi - aanhouden - afgelegen - rijden
**snel** - hoger - vlug - financieel - dor - diner - traag - hevig - bron - rap - bunker
**geit** - aanloop - bok - aanrecht - kaas - bevel - kort - dier - borst - boerderij
**parlement** - minister - atlas - bezoeker - regering - gedragen - ellende - politiek - afzender
**lijm** - hecht - kleven - haten
**schorpioen** - giftig - minuut - groeten - doodstil - verschil - raam - gif - colonne - concert - woestijn
**zondag** - haak - rust - attractie - bankier - anders - vrij - stil - bang - weekend - prachtig
**rits** - open - grazen
**lucifer** - vuur - feit - zwavel - baat - brand - aandeel - inzicht - borrel - afdalen - sigaret
**politieagent** - uniform - geheim - boete - bakermat - halen - bijzijn - docent
**doorzichtig** - slaaf
**mooi** - beurt - links - soort - mantel - borg - lelijk - uitspraak - rand - dreiging - knap
**tandpasta** - kost - gewoon - college - tandenborstel - flauw - poetsen - jong - bedwang - munt - zorgen
**praten** - boon - farao - gesprek - nauw - extra - echt - mond - aura - badjas - babbelen
**geest** - spook - mening - ziel - bordeaux - vals
**baby** - schattig - even - landen - apotheek - armslag - huilen - lepel - commandant - terecht - luier
**tong** - smaak - flat
**dak** - schoorsteen - opkomen
**goud** - zilver - alinea - hou - tikken - gelukkig - rijk - dekker - reactie - ervaring - duur
**kabouter** - paddestoel - begin - heersen - muts - belevenis - lied - amateur - tuin - grijpen - dwerg
**team** - cognac - sport - buigen - plek - groep - afspraakje - samen - abces - taal - ploeg
**telefoon** - bellen - lezer - nummer - bons
